# G1/S Boundary Activates Interferon and Inflammatory Response Genes

**DOI:** 10.1101/2023.08.24.554683

**Authors:** Gözde Büyükkahraman, Tae Hoon Kim

## Abstract

Interferons (IFNs) have various roles in antiviral immunity, including curbing the immune system to prevent tissue damage and stimulating adaptive immunity. Due to its protective and destructive properties, IFN expression is tightly regulated. In contrast to its tight regulatory control, IFN expression is highly heterogeneous across many cell types upon pathogenic stimulus. The basis for this heterogenous IFN expression remains incompletely understood. Using single cell RNA-sequencing upon viral infection, we found that interferon expression is upregulated specifically in the late G1 phase of the cell cycle, and cell synchronization at the G1/S boundary boosts interferon expression. Furthermore, cell cycle arrest without any additional stimulus is sufficient to upregulate interferons and hundreds of other inflammatory response genes. Interferon upregulation at the G1/S boundary is cell type specific and not observed in non-immune cell types. Finally, we use ATAC-seq to identify potential transcription factors orchestrating this response. Together, these results uncover the cell cycle as a critical regulator of IFN expression in immune cells.

## Introduction

The type I interferon (IFN) system acts as a first line of defense before the more effective adaptive immune system can mount a response. Upon infection, viral RNA is detected in the cytoplasm of the host cell by RNA helicases (Kato, Takeuchi et al. 2006). When these RNA helicases bind to viral RNA, they form heterodimers, go through a conformational change, and activate transcription factors (TFs) IFN regulatory factors 3, 7 (IRF3 and IRF7), and NF-κB. These TFs then translocate into the nucleus to drive the expression of *IFNB1*, a prototypical type I IFN cytokine gene (Fitzgerald, McWhirter et al. 2003, Sharma, tenOever et al. 2003, tenOever, Sharma et al. 2004). After secretion, IFN-β cytokine binds to interferon alpha receptor (IFNAR) to induce expression of interferon stimulated genes (ISGs) and ensure antimicrobial defense. Due to its centrality in antimicrobial response, regulation of IFN expression is tightly regulated, and abnormalities in this regulation are linked to various diseases (Ivashkiv and Donlin 2014, McNab, Mayer-Barber et al. 2015).

Dysregulation of type I IFNs have been extensively investigated in many diseases. These involve COVID-19 patients with poor prognosis (Lowery, Sariol et al. 2021, Stephenson, Reynolds et al. 2021), autoimmune diseases with type I IFN overexpression (Uggenti, Lepelley et al. 2019), and sub-optimal type I IFN expression that leads to reduced protection against pathogens (Hadjadj, Yatim et al. 2020). These observations show that proper type I IFN regulation requires an intricate balance that is not always properly achieved. Molecular studies also revealed that type I IFN levels can be highly variable across individual cells. IFN expression heterogeneity can be due to variations in viral infection, host state, spatiotemporal heterogeneity of the cells, and complex interactions between these parameters (Van Eyndhoven, Singh et al. 2021). For example, viruses rely on host factors for effective infection and can cause heterogeneous expression of type I IFNs. The latest studies examined gene expression heterogeneity across infected cells at a single-cell resolution and concluded that IFN expression heterogeneity may come from the differences in the virus’ ability to induce an immune response (Lauring, Frydman et al. 2013, Stern, Bianco et al. 2014). However, other studies argued that the cause of IFN expression heterogeneity is not the viral abundance within the cell but rather the internal factors such as the host cell state (Rand, Rinas et al. 2012, O’Neal, Upadhyay et al. 2019).

Several studies have suggested that the cell cycle phase of the host cell at the time of infection may determine whether the infected cell will express IFN-β or not (Lee and Rozee 1970, Bressy, Droby et al. 2019). In recent studies, involvement of cell cycle on type I IFN expression regulation was investigated; however, there is no consensus on what cell cycle phase or the factors involved in this expression (Lee and Rozee 1970, Genin, Cuvelier et al. 2015, Cingoz and Goff 2018). In addition to type I IFNs, expression of other cytokines was also found to be variable across responsive cells (Calado, Paixao et al. 2006, Mariani, Schulz et al. 2010, Bass, Wong et al. 2021). Variability in the cytokine expression across a seemingly homogeneous cell group must be studied at a single cell resolution as the transcriptional differences of a few responding cells can be masked due to averaging of the expression levels in the bulk studies (Satija and Shalek 2014, Drayman, Patel et al. 2019).

In this study, we take advantage of the cellular resolution of the single cell RNA sequencing to understand the extent of IFN expression heterogeneity in GM12878 cells. We observed the most significant IFN induction in cells of late G1 phase of the cell cycle. Surprisingly, arresting cells at G1/S, without any pathogenic stimuli, induced a strong immune response by upregulating hundreds of cytokine genes in addition to IFNs. G1/S induction of immune response was cell type specific, observed only in immune cells. Upon G1/S phase arrest, thousands of regulatory sites near immune response related genes became more accessible. Combining bulk RNA sequencing and bulk ATAC sequencing revealed a list of known immune response TFs which may drive G1/S arrest-mediated immune response. Our results indicate that immune cells in late G1 phase become more competent to induce immune response through accumulated expression of TFs and associated changes in chromatin accessibility of relevant immune response genes.

## Results

### scRNA-seq Reveals IFN Upregulation in Late G1

To investigate the cause of IFN expression heterogeneity in an unbiased manner, we performed single cell RNA sequencing (scRNA-seq) after infection of a B lymphoblastoid cell line (GM12878) with Sendai virus (SeV). SeV is a single-stranded RNA virus which has been used to study anti-viral response in a wide range of hosts (Strahle, Garcin et al. 2003). After quality control, we detected ∼11,000 cells. We labeled cells with at least one copy of the mRNA for SeV genes -*M*, *L*, *N*, *P*, *HN*, and *F*-as “Viral mRNA+”, and the rest as “Viral mRNA-”. After dimensionality reduction, we observed that the cells were clustered into two major groups, which are largely driven by the viral mRNA presence **(Figure 1a)**.

**Figure 1:**
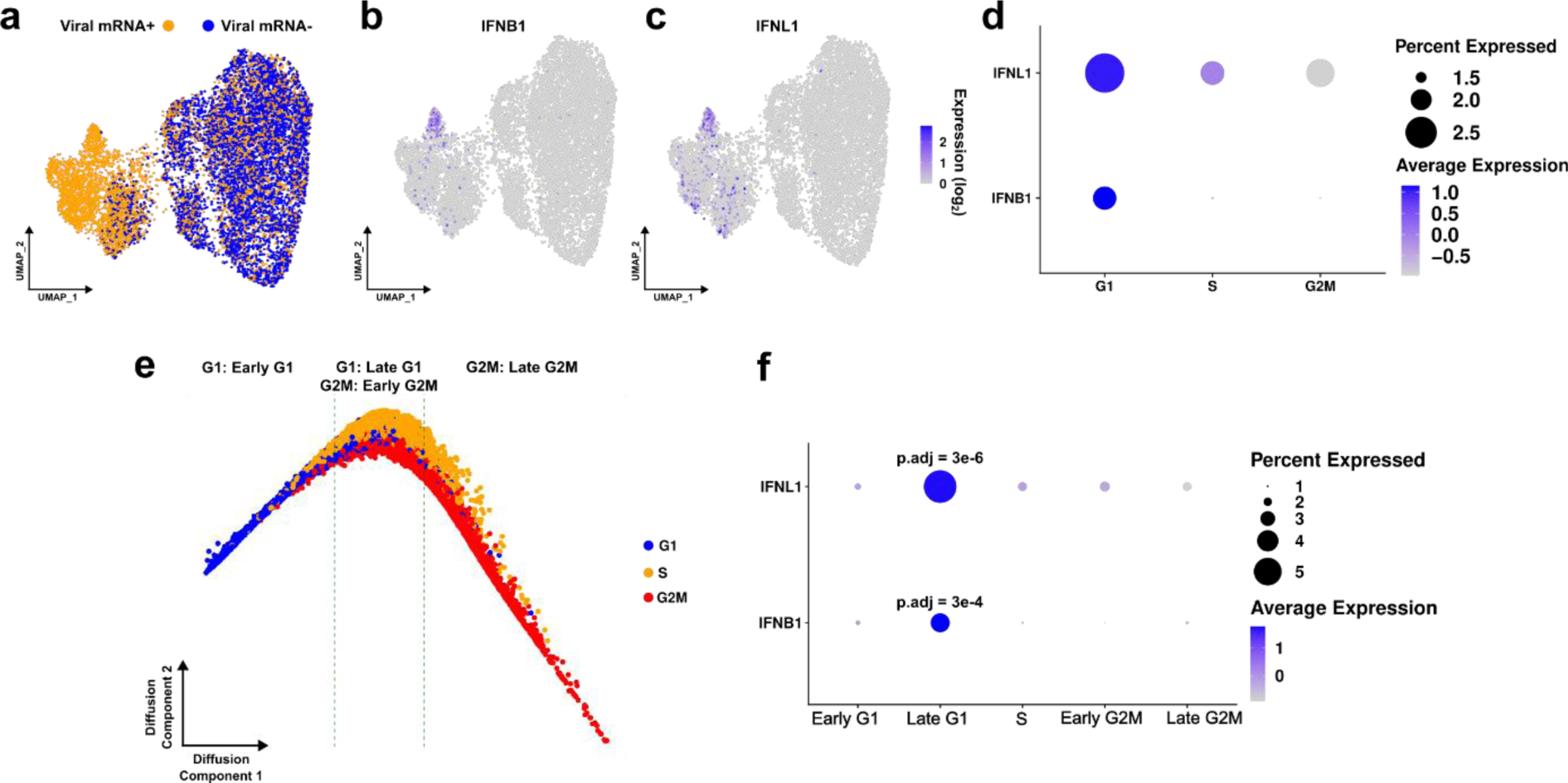
scRNA-seq of SeV infected GM12878 (GM) cells. **(a)** UMAP of infected single cells labeled according to viral mRNA presence. The cells with >= 1 read mapping to any one of the viral genes (*HN*, *L*, *P*, *M*, *N*, and *F*) were grouped as Viral mRNA+. Viral mRNA-cells do have any reads of viral genes. Viral mRNA+ cells are shown in yellow whereas Viral mRNA-cells are shown in blue. **(b)** *IFNB1* expressing cells across the same UMAP as (a). Dark blue indicates high expression whereas light blue indicates low expression. **(c)** *IFNL1* expressing cells across the same UMAP as (a). Dark blue indicates high expression whereas light blue indicates low expression. **(d)** Bubble chart showing the expression of *IFNB1* and *IFNL1* across cell cycle phases. Bigger dots indicate higher number of cells expressing the gene. Dark blue indicates high expression whereas light blue indicates low expression. **(e)** Pseudotime trajectory of single cells according to their cell cycle scoring. Blue indicates the cells in G1 phase, yellow indicates the cells in S phase, and red indicates the cells in G2M phase. The cells were grouped into 5 subgroups of cell cycle phases: Early G1, late G1, S, early G2M, and late G2M. **(f)** Bubble chart showing the expression of *IFNB1* and *IFNL1* across subgroups of cell cycle phases. Bigger dots indicate higher number of cells expressing the gene. Dark blue indicates high expression whereas light blue indicates low expression. Adjusted p-values were obtained by Wilcoxon rank sum test and Bonferroni correction.

We noticed that type I and type III IFNs (hereby referred to as IFNs for simplicity) were heterogeneous among the viral mRNA+ population, with some cells displaying high amounts of *IFNB1* and *IFNL1* (two of the most upregulated IFNs upon infection in GM cells) expression **(Figure 1Figure 1b-c)**. This IFN expression heterogeneity is in line with the previous reports (Cai, Zhou et al. 2017). To investigate how the host cell state might be affecting the IFN expression, we focused on the cell cycle phase, which has been previously implicated in immune-mediated responses (Genin, Cuvelier et al. 2015, Bressy, Droby et al. 2019, Daniel, Belk et al. 2023). We grouped cells according to their cell cycle phases using gene expression differences that mark S and G2/M phases (Hao, Hao et al. 2021). This resulted in G1, S, and G2/M categories and revealed higher *IFNB1* and *IFNL1* levels in the G1 phase, although this was not a statistically significant difference **(Figure 1d)**. Since G1 is a long cell cycle phase that lasts about 12h (Chen 1992, Lee, Goolsby et al. 1994), we then sought to further resolve cells within G1 phase using pseudotime (Hao, Hao et al. 2021) and grouped them into 5 clusters: early G1, late G1, S, early G2M, and late G2M **(Figure 1e)**. Among these clusters, *IFNB1* and *IFNL1* expression levels were significantly upregulated in late G1/S phase cells **(Figure 1f)**. These results indicate that the cell cycle state of the host might be an important determinant of IFN expression upon viral infection.

### G1/S Arrest Coupled with Infection Boosts IFN Expression

To understand which cell cycle phase is the most permissive to IFN expression, GM12878 cells were treated with cell cycle phase blockers and infected with SeV. To ensure the cells were arrested at the indicated cell cycle phase, we treated the cells with small molecule drugs aphidicolin and nocodazole at different time points. Aphidicolin is a small molecule that reversibly binds to or near the nucleotide-binding site of DNA polymerase and inhibits DNA replication, thereby pausing the cells at the G1/S boundary (Krokan, Wist et al. 1981). Nocodazole is another small molecule that arrests the cell at the G2/M boundary by preventing the formation of microtubules (Samson, Donoso et al. 1979). Using flow cytometry, we observed that treatment of GM12878 with aphidicolin arrested the cells at the G1/S boundary, and treatment with nocodazole arrested the cells at the G2/M boundary, as expected **(Supplementary Figure 1)**.

After cell cycle phase synchronization, we infected the cells with SeV and found higher expression of *IFNB1* in G1/S arrested cells compared to the control, whereas G2/M arrested cells displayed similar *IFNB1* levels to the control **(Figure 2a)**. These findings further support that IFN transcript levels increase significantly at the G1/S boundary.

**Figure 2:**
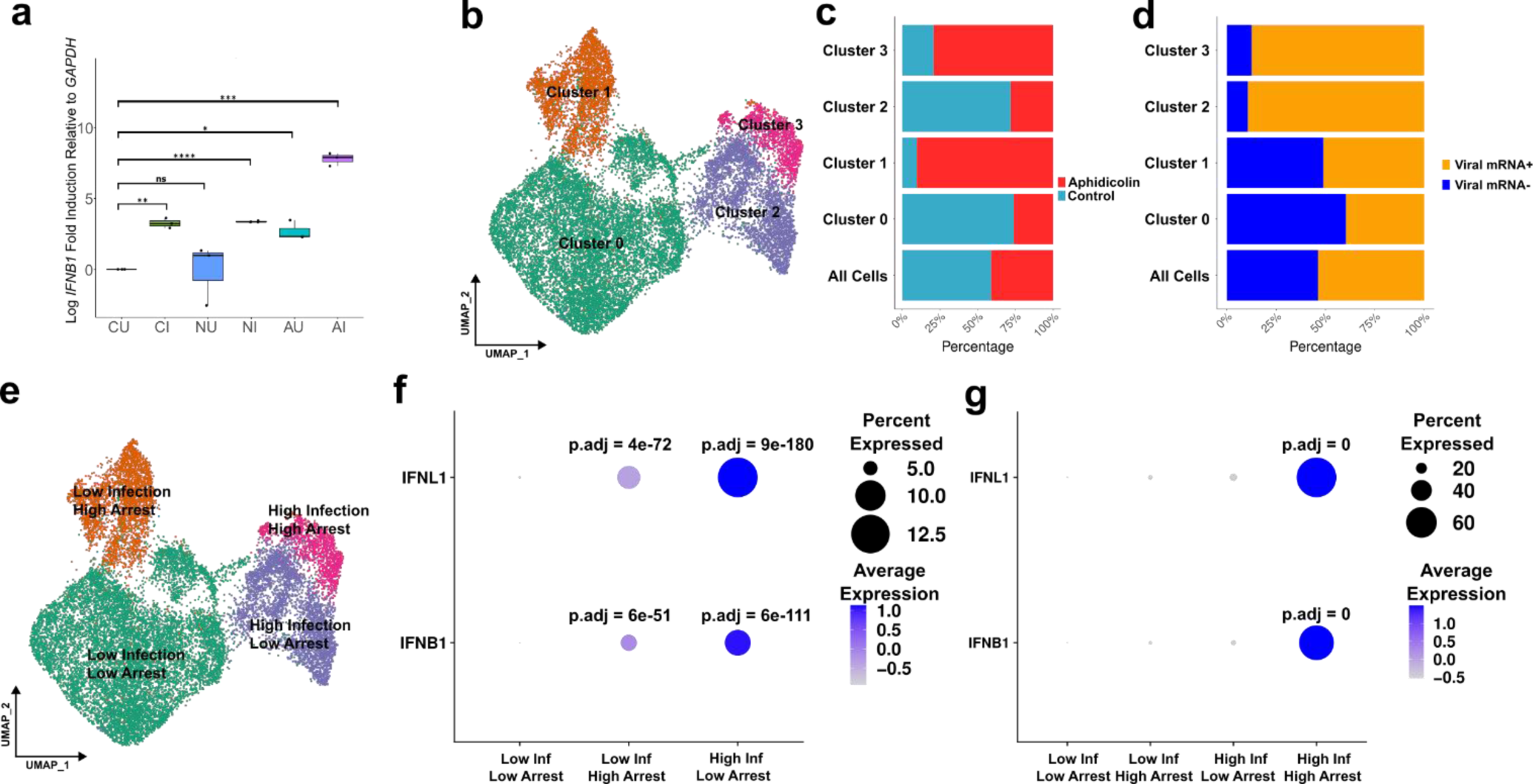
Effect of cell cycle arrest on IFN expression. **(a)** Log transformed fold inductions of *IFNB1* relative to *GAPDH* after infection and/or nocodazole treatment, and aphidicolin treatment. C represents control (DMSO treated), N represents nocodazole treated samples, and A represents aphidicolin treated samples. I represents the SeV infected samples and U represents the uninfected controls. Statistical significance was determined by t-test. ns indicates p value > 0.05, * indicates p value < 0.05, ** indicates p value < 0.01, *** indicates p value < 0.001, **** indicates p value < 0.0001, n=3. **(b)** UMAP of batch corrected and clustered single cells. **(c)** The proportion of control cells (DMSO treated) and aphidicolin treated cells across clusters. Red indicates aphidicolin treated cells whereas blue indicates control cells. **(d)** The proportion of cells with the viral mRNA across clusters. The cells were grouped in terms of presence of viral mRNA as mentioned in Figure 1. **(e)** Relabeled clusters in the same UMAP as (a). Clusters were relabeled according to their sample and viral mRNA cell proportions in (b) and (c). **(f)** Bubble chart showing the expression of *IFNB1* and *IFNL1* across clusters. Bigger dots indicate higher number of cells expressing the gene. Dark blue indicates high expression whereas light blue indicates low expression. **(g)** Bubble chart showing the expression of *IFNB1* and *IFNL1* across clusters. Bigger dots indicate higher number of cells expressing the gene. Dark blue indicates high expression whereas light blue indicates low expression. Adjusted p-values were obtained by Wilcoxon rank sum test and Bonferroni correction.

To understand the induction of IFNs at the G1/S boundary at a cellular resolution, we performed scRNA-seq on cells after G1/S arrest and viral infection. After quality control filtering, we detected ∼7500 cells and a similar number of reads to the control group – infected with the virus but not arrested – **(Supplementary Figure 2a-b)**. After removing the batch effect between the two conditions (G1/S arrested-infected and infected conditions), we identified 4 major clusters **(Supplementary Figure 2c-d, Figure 2b)**. We noticed that the cells were clustered based on their treatment (control or G1/S arrested) and infection (uninfected or infected) levels **(Figure 2c-d, Supplementary Figure 2e-f)**. We labeled the clusters accordingly as low/high arrested and low/high infected **(Figure 2e)**. Clusters mainly characterized by the arrest or infection both displayed significantly higher IFN levels compared to the cluster that had low levels of both **(Figure 2f)**. Strikingly, IFN levels were boosted in the cluster with both high proportions of arrested and infected cells **(Figure 2g)**. These findings indicate that both G1/S boundary and viral infection are critical regulators of IFN expression.

### G1/S Arrest Alone is Sufficient to Induce Immune Response

IFNs are expressed and secreted in response to a pathogenic stimulus as the first line of defense. Therefore, studies focusing on the regulation of IFNs typically utilize an external stimulant (e.g., virus infection) to induce IFN expression (Zhao, Zhang et al. 2012). To understand whether the G1/S boundary can induce IFN expression in the absence of viral infection, we performed bulk RNA sequencing for 4 conditions: control uninfected (CU), control infected (CI), G1/S arrested and uninfected (AU), and G1/S arrested and infected (AI). To assess if cell synchronization at the G1/S phase was effective, we plotted the expression levels of S phase marker genes across treatment groups. As expected, arrested cells had higher expression levels for S markers *CCNE2*, *E2F8*, *CDC6*, and *CDC45* **(Supplementary Figure 3a)**. Overall, we found that most of the variability in the data could be explained by G1/S arrest (PC1: 61%) and viral infection (PC2: 25%) **(Figure 3a)**. IFN levels were significantly upregulated in CI compared to CU, as expected **(Figure 3b-c, Supplementary Table 1)**. Remarkably, G1/S arrest alone was sufficient to cause a significant induction of *IFNB1* (∼42 fold) and *IFNL1* (∼13 fold) **(Figure 3b-c, Supplementary Table 1)**. The induction level increased dramatically when cells were infected and arrested **(Figure 3b-c)**. These results indicate that IFN expression can be robustly induced by G1/S arrest in the absence of viral infection.

**Figure 3:**
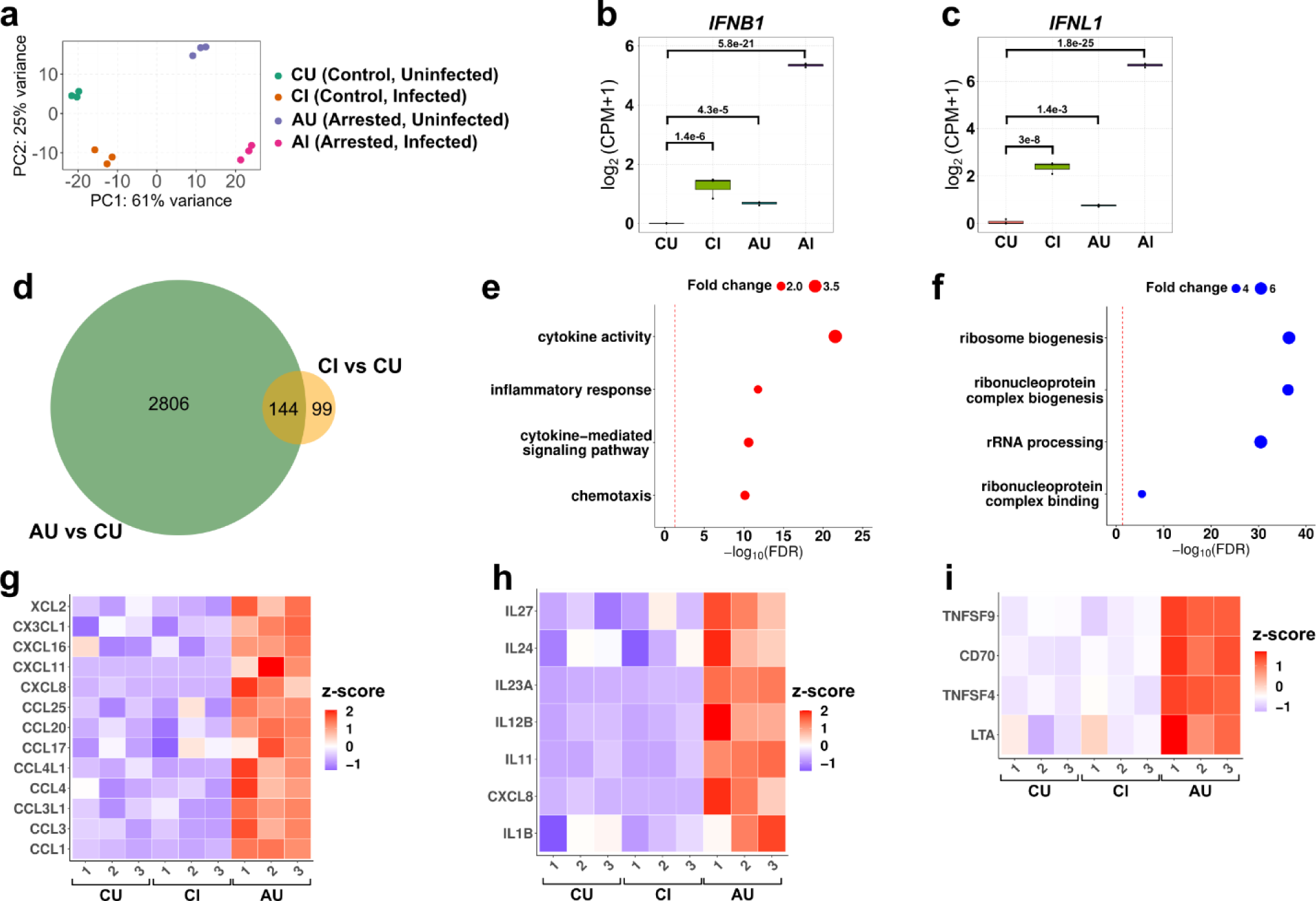
RNA-seq results of GM12878 (GM) cells after G1/S arrest and/or viral infection. GM cells were treated with DMSO (CU; control uninfected), SeV (CI; control infected), aphidicolin (AU; arrested uninfected), and aphidicolin and SeV (AI; arrested uninfected). **(a)** PCA plot showing the gene expression data variability across samples. Note that groups are well separated in PC1 and PC2 dimensions. **(b-c)** The box plot of log normalized **(b)** *IFNB1* and **(c)** *IFNL1* counts across samples. Statistical significance was determined by Wald test. **(d)** Venn diagram showing the number of common differentially expressed genes in AU-CU comparison and CI-CU comparison. **(e-f)** Bubble charts of significantly enriched gene ontology (GO) terms among the **(e)** upregulated differentially expressed genes (DEGs) and **(f)** downregulated DEGs in AU samples compared to CU samples. Statistical significance was determined by Fisher’s exact test. **(g-i)** Heatmaps of chemokines (g), interleukins (h) and tumor necrosis factors (i) across CU, CI, and AU samples. Numbers denote biological replicates. Gene expression values were normalized and z-transformed across all samples.

We then focused on the global gene expression changes upon G1/S arrest and viral infection. Differential gene expression analysis showed that the number of significantly upregulated genes was more than the significantly downregulated genes upon G1/S arrest and/or upon infection **(Supplementary Figure 3b)**. Combined arrest and infection resulted in more differential gene expression changes than either arrest or infection alone **(Supplementary Figure 3b)**. Most of the genes which were induced by infection alone (CI) or by G1/S arrest alone (AU) were also induced by both treatments combined, indicating that arrest and infection largely result in the gene expression changes in a similar direction **(Supplementary Figure 3c-d)**. We also found a large overlap of differentially expressed genes between AU-CU and CI-CU comparisons that support the conclusion that the infection and arrest lead to differential expression of a similar set of genes **(Figure 3d)**.

Gene ontology enrichment of genes upregulated in arrested cells (AU > CU) revealed inflammatory response, cytokine, and chemotaxis-related genes as the top enrichments **(Figure 3e and Supplementary Table 2)**. In contrast, downregulated genes (AU < CU) were enriched in ribosome biogenesis, ribonucleoprotein binding, and rRNA process-related genes **(Figure 3f and Supplementary Table 2)**. Despite the large overlap between the genes differentially expressed upon G1/S arrest and infection, we detected ∼2800 genes that were significantly induced by arrest but not by infection **(Figure 3d)**. Interestingly, these genes included many cytokines, namely interleukins, chemokines, and TNFs that were not upregulated upon infection **(Figure 3g-i)**. This contrasts with the IFN expression that is similarly upregulated by infection and by G1/S arrest. These results suggest that G1/S arrest induces a broader immune response than viral infection alone, and this response involves many cytokines in addition to IFNs.

### G1/S Arrest-mediated IFN Induction is Cell Type Specific

We analyzed other cell lines to determine whether this effect is general and applicable to other cell types. We analyzed Ramos, a non-EBV-infected B cell line, THP-1, a monocyte cell line, and Jurkat, a T cell line **(Supplementary Figure 4a-d, e-f, i-j)**. All cell lines showed significantly increased *IFNB1* mRNA levels during the G1/S phase of the cell cycle, and infection can further elevate their expression, similar to GM12878 **(Figure 4a-c)**. Given that the G1/S phase activates the IFN expression in immune cell lines, we asked whether non-immune cells could exhibit this strong expression pattern. We similarly treated IMR90 cells with aphidicolin and SeV. IMR90 did not show an increase in *IFNB1* mRNA levels upon G1/S arrest **(Figure 4d and Supplementary Figure 4g-h)**. These results suggest that G1/S arrest-mediated upregulation of IFNs is cell-type specific.

**Figure 4:**
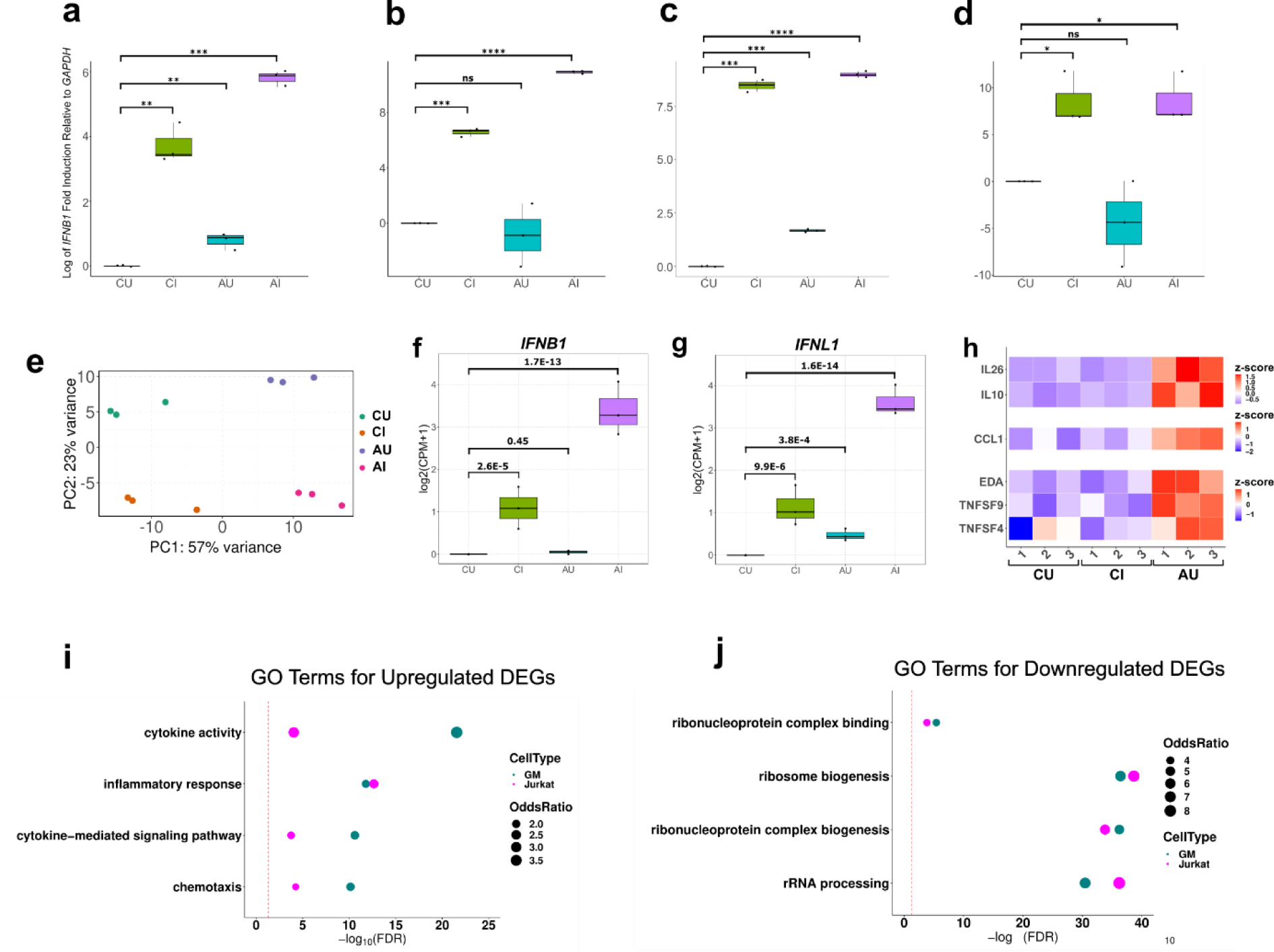
G1/S arrest mediated IFN upregulation across multiple cell types. **(a-d)** Log transformed fold inductions of *IFNB1* levels across treatment conditions DMSO (CU), SeV (CI), aphidicolin (AU), and aphidicolin and SeV (AI) in cells **(a)** Jurkat, **(b)** Ramos, **(c)** THP-1, and **(d)** IMR90. Statistical significance was determined by t-test. ns indicates p value > 0.05, * indicates p value < 0.05, ** indicates p value < 0.01, *** indicates p value < 0.001, **** indicates p value < 0.0001, n=3. **(e)** PCA plot showing the gene expression data variability across samples. Note that groups are well separated in PC1 and PC2 dimensions. **(f-g)** The box plot of log normalized **(f)** *IFNB1* and **(g)** *IFNL1* counts across samples. Statistical significance was determined by Wald test. **(h)** Heatmaps of interleukins (*IL26* and *IL10*), chemokines (*CCL1*) and tumor necrosis factors (*EDA*, *TNFSF9*, and *TNFSF4*) across CU, CI and AU samples. Numbers denote biological replicates. Gene expression values were normalized and z-transformed across all samples. **(i-j)** Bubble charts of significantly enriched gene ontology (GO) terms among the **(i)** upregulated differentially expressed genes (DEGs) and **(j)** downregulated DEGs in AU samples compared to CU samples. Statistical significance was determined by Fisher’s exact test.

To understand the global transcriptomic differences or similarities between B and T cell lines upon G1/S arrest, we performed bulk RNA sequencing with Jurkat cells using 4 treatment groups like GM12878 **(Figure 4e)**. S-phase marker genes were similarly upregulated in Jurkat upon G1/S arrest **(Supplementary Figure 3e and Supplementary Table 3)**. As seen in GM12878, we detected more upregulations than downregulations upon G1/S arrest **(Supplementary Figure 3f)**, and we detected more differential gene expression in combined arrest and infection than either arrest or infection alone **(Supplementary Figure 3f)**. Similarly, genes upregulated in only arrest or only infection were largely detected in combined arrest and infection **(Supplementary Figure 3g-h).** Thus, Jurkat displayed many similar expression patterns seen in GM12878 cell line. However, unlike GM12878, Jurkat did not show significant induction in *IFNB1* upon G1/S arrest alone, although *IFNL1* induction was significant **(Figure 4f-g and Supplementary Table 3)**. Nevertheless, the induction of both cytokines was similarly boosted upon combined G1/S arrest and viral infection, indicating the expression-inducing effect of G1/S arrest on both *IFNB1* and *IFNL1* **(Figure 4f-g and Supplementary Table 3)**. Inflammatory response-related genes were enriched in the upregulated set of genes. Ribosome biogenesis related genes were enriched in downregulated genes **(Figure 4i-j and Supplementary Table 4)**. However, compared to GM12878, we detected lower enrichment in cytokine and chemokine-related genes **(Figure 4h)**. Indeed, the number of cytokines that were upregulated in G1/S compared to the controls was lower than in GM12878. They were similarly only upregulated by G1/S arrest and not by infection **(Figure 4h)**. Despite the differences, G1/S arrest-induced genes were significantly common between these cell lines **(Supplementary Figure 3i)**. These results support similar inflammatory responses between B and T cell lines but also suggest cell type specific differences notably in the regulation of cytokines.

### ATAC-seq Reveals G1/S Arrest Mediated-Putative TF Regulators

Gene expression changes are typically coupled with chromatin accessibility changes. To understand the accessibility changes associated with G1/S arrest-mediated IFN and inflammatory response upregulation, we performed Assay for Transposase-Accessible Chromatin sequencing (ATAC seq) of the same treatment groups (CU, CI, AU, and AI) in GM12878. We first identified peaks that are consistently observed across the biological replicates for each of the 4 treatment groups. Distribution of these peaks across the annotated promoters, introns, exons, UTRs, and intergenic regions showed an expected distribution (Yan, Powell et al. 2020) **(Supplemental Figure 5a-d)**. We then merged these peaks, resulting in ∼200,000 peaks that were used as a common set of peaks to compare the genomic accessibility across treatments. Similar to bulk RNA-seq, most of the variability within the data could be explained by G1/S arrest (PC1: 66% of the variance) **(Supplementary Figure 5e)**.

To reveal accessibility differences between G1/S arrested and control groups, we identified differentially accessible regions (DARs) on the common set of peaks. This revealed thousands of chromatin accessibility changes upon G1/S arrest **(Supplementary Figure 5f, Supplementary Table 5)**. DARs upon G1/S arrest and upon combined arrest and infection also displayed a large overlap, as expected based on the bulk RNA-seq results **(Supplementary Figure 5g)**. We then performed gene ontology enrichment on the chromatin accessibility changes upon G1/S arrest using GREAT (McLean, Bristor et al. 2010). Top enrichments for the significantly open DARs in G1/S arrest included immune system process and inflammatory response, as were seen in the bulk RNA-seq results **(Figure 5a, Supplementary Table 6)**. In contrast, enrichments for the significantly closed DARs included neuronal and cytoskeletal-related terms **(Figure 5b, Supplementary Table 6)**. To identify potential gene regulatory elements (GREs) around interferon genes, we scanned IFN loci for DARs. In both the locus containing *IFNL1-4* and *IFNB1*, we identified GREs that were significantly more accessible upon G1/S arrest **(Figure 5c-d)**. Two of these GREs were also previously identified using H3K27ac and H3K4me1 chromatin immunoprecipitation analyses (Zhang, Lee et al. 2020), indicating that these are active regulatory elements prior to G1/S arrest or infection. Notably, one of these regions (L2 enhancer) has been previously validated to induce *IFNB1* expression upon viral stimulation **(Figure 5d)** (Banerjee, Kim et al. 2014). These results indicate that without any pathogenic stimulus, G1/S arrest increases chromatin accessibility near IFNs and other immune response genes.

**Figure 5:**
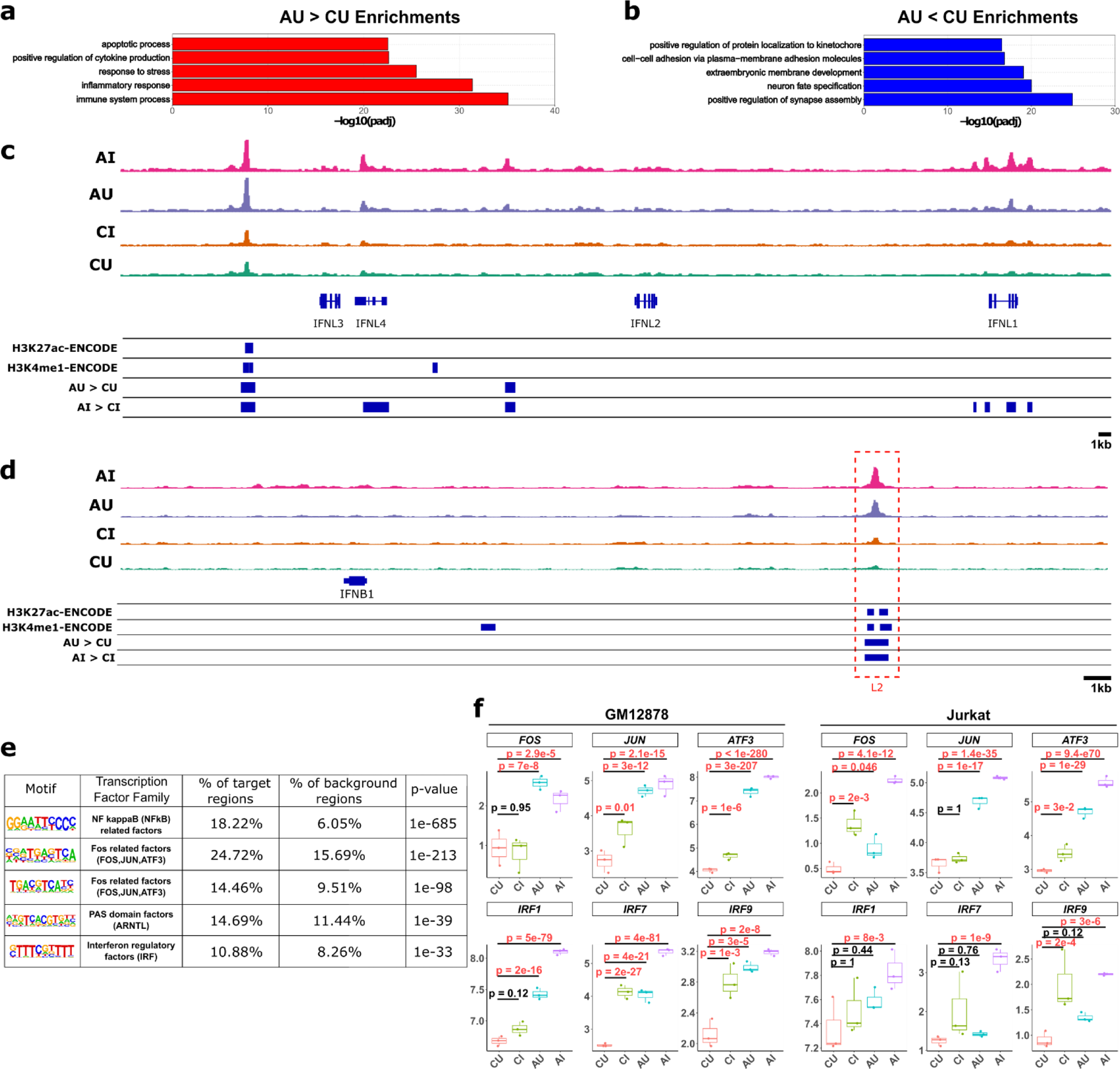
ATAC-seq reveals gene regulatory elements and potential TF regulators orchestrating immune response upon G1/S arrest. **(a-b)** Bar plots of significantly enriched gene ontology (GO) terms among the **(a)** differentially open genomic regions in GM cells treated with aphidicolin (AU) compared to control (CU) and **(b)** differentially close genomic regions in AU compared to CU. Statistical significance was determined by Fisher’s exact test. **(c-d)** Normalized peaks detected across CU, CI, AU, and AI samples along with ENCODE H3K27ac and H3K4me1 data and differentially accessible regions in AU-CU and AI-CI comparison. These peaks are shown for (c) *IFNL1-4* loci and (d) *IFNB1* loci. Note that L2, known enhancer of *IFNB1*, is highlighted in red rectangle. **(e)** Motif enrichment analysis of differentially open chromatin regions in AU-CU comparison. **(f)** Transcription factor expression levels in GM and Jurkat bulk RNA-seq data across treatment conditions. Statistical significance was determined by Wald test.

To understand the potential transcriptional factors (TFs) that regulate chromatin accessibility, we performed TF motif enrichment analysis among DARs upon G1/S arrest. We found NF-κB related factors, AP-1 family transcription factors, PAS domain-containing factors, and IRF factors to be significantly enriched in DARs compared to the background (**Figure 5e and Supplementary Table 7**). Among these TF families AP-1 related factors and IRFs are involved in regulating the immune system (Wagner and Eferl 2005, Nehyba, Hrdlickova et al. 2009, Li, Spolski et al. 2012, Atsaves, Leventaki et al. 2019). We found that *FOS*, *JUN*, *ATF3*, *IRF1*, *IRF9,* and *IRF7* were significantly upregulated upon G1/S arrest in both GM12878 and Jurkat cells **(Figure 5f and Supplementary Tables 1-3)** indicating that these TFs may be involved in the induction of immune response upon G1/S arrest by changing the chromatin accessibility. Taken together, these results reveal potential TFs orchestrating gene regulatory response upon cell cycle arrest at G1/S boundary.

## Discussion

In this study, we focused on the causes of IFN expression heterogeneity. Our single cell RNA-sequencing analysis shows that late G1 phase cells produce more IFN compared to other cell cycle phases upon infection. Strikingly, G1/S phase arrest without any viral infection also leads to robust activation of IFN cytokines, and G1/S arrest coupled with viral infection leads to enhanced IFN expression. We also show that G1/S arrest alone leads to upregulation of many cytokines, including chemokines, interleukins, and TNFs, indicating a stronger immune response in G1/S arrest. This observation suggests that G1/S transition can elicit expression of a broader set of immune response genes compared to simple viral infection. Combination of both infection and appropriate cell cycle phase can drive stronger IFN expression. Interestingly, IFN upregulation at G1/S arrest is cell type specific and observed in immune cell lines, but not in non-immune cells that we have examined. Through ATAC-seq, we identified relevant cis-regulatory elements that become more accessible upon G1/S arrest, including known and potential enhancers around IFN genes. Finally, motif enrichment analyses on these elements propose NF-κB, AP-1, and IRF family TFs as potential orchestrators of gene regulatory changes upon G1/S arrest.

IFN is the first cytokine to be expressed upon viral infection. Despite its importance in inducing antiviral state, IFN expression was observed in only 10-30% of cells in a population which was uniformly infected by virus (Zhao, Zhang et al. 2012). As a potential source of IFN expression heterogeneity, prior studies focused on the differences in the ability of a virus to induce host immune response (Killip, Young et al. 2011, Rand, Rinas et al. 2012, Zhao, Zhang et al. 2012, Patil, Fribourg et al. 2015, Russell, Elshina et al. 2019). These studies reported that the variability in the virus can contribute however it does not fully explain the heterogeneity in IFN expression (Rand, Rinas et al. 2012, O’Neal, Upadhyay et al. 2019, Russell, Elshina et al. 2019). Here, we report significant induction of IFN expression upon G1/S arrest alone without any virus infection, suggesting that this source of IFN expression heterogeneity is intrinsic to the cell state **(Figure 3b-c and Figure 4f-g)**. We also observed that G1/S arrest combined with infection results in an even stronger expression of immune related genes. Therefore, the differences in the intrinsic cell cycle phases and infecting virus can combine to drive highly variable expression of IFN expression. This coupling suggests that interferon and immune response in these cells is tightly integrated with other cellular signals that promote cell cycle progression and proliferation.

Cells can respond to a plethora of stimuli to express immune genes: virus, bacteria, pathogen-mimicking particles, poly (I:C), LPS, IFN-α, and IFN-β (Marshall, Warrington et al. 2018). Both pathogens and IFNs have been shown to arrest the host cell cycle phase and influence the immune response (Xaus, Cardo et al. 1999, Yuan, Shan et al. 2005, Dove, Brooks et al. 2006, Ambjorn, Ejlerskov et al. 2013, Maeda, Wada et al. 2014, Brenner, Schorg et al. 2020). Others have taken a more direct approach and tracked cell cycle phases of live macrophages prior to LPS induction followed by single cell RNA sequencing. They found that S phase cells showed less immune response than other phases, concluding that the cell cycle phase affects the immune response to stimulus but not the vice versa (Chen, Guillaume-Gentil et al. 2022). Here, by combining pseudotime analysis of scRNA-seq data and arresting cell cycle progression, we provide further support that cell cycle phase of the host affects the immune response. Moreover, we provide the first direct evidence that cell cycle arrest at the G1/S boundary is sufficient to drive immune response even in the absence of viral infection. Strikingly, this response is stronger than the immune response produced by viral infection alone in these cells **(Figure 3g-i and Figure 4h)**. Our results highlight the importance of the cell cycle in dysregulated immune response, especially in autoimmune diseases. Increased proliferation of these immune cells alone can drive the production of these inflammatory cytokines.

Type I IFNs are universally expressed in almost all cell types to induce an antiviral state throughout the organism (McNab, Mayer-Barber et al. 2015). Effects of the cell cycle in type I IFN expression have been previously investigated across various cell types: G2/M phase was found to suppress IFNs and enhance viral replication in human pancreatic cancer cell lines (Bressy, Droby et al. 2019). A previous report found that CDK1/2 activity essential for mitosis is required for type I IFN translation in human monocytes and fibroblasts (Cingoz and Goff 2018). There is no consensus on the specific cell cycle phase that may be involved in IFN expression heterogeneity, and the underlying mechanisms of this heterogeneity remain far from being resolved. In our present work, we report that G1/S arrest alone can induce IFNs in B and T cells, immune cells, but not in other cell types, non-immune cells **(Figure 4a-d)**. Interestingly fibroblasts did not exhibit cell cycle dependence of IFN expression. **(Figure 4d)**. These results suggest that the different cell types can exhibit distinct underlying mechanisms for IFN expression heterogeneity.

How cell type specific molecular mechanisms induced by G1/S arrest specifically upregulate immune response genes remains to be determined. One hypothesis for the underlying mechanism involves double-strand breaks (DSBs) that can be induced by G1/S arrest (Mazouzi, Stukalov et al. 2016, Ma, Feng et al. 2021). A previous study done in neurons found that DSBs alone can induce the expression of genes immediately upregulated by neuronal activity. (Madabhushi, Gao et al. 2015). Here we show that another inducible gene regulation model, IFN response to pathogenic stimulus, can be activated by G1/S arrest alone in the absence of viral infection. Therefore, G1/S arrest of B and T cell lines may induce a DSB and DNA damage repair mediated IFN activation. Interestingly, we found that DNA damage response gene *E2F7*, single-strand break repair scaffold protein, *XRCC1*, and a CDK inhibitor, *CDKN1A*, which is driven by replication stress-induced DNA damage, are all upregulated upon G1/S arrest **(Supplementary Tables 1 and 3)** (Hume, Dianov et al. 2020). Other studies support the role of DSB and DNA repair in modulating immune response. For example, inhibition of topoisomerase, a protein involved in DNA replication and repair, was found to reduce pro-inflammatory cytokine expression and to protect the organism from sepsis (Rialdi, Campisi et al. 2016, Ho, Mok et al. 2021, Zhang, Zhao et al. 2021). Taken together with the existing literature, upregulation of immune response and DNA repair genes by G1/S arrest in the absence of viral stimulus indicates a close interplay between cell cycle, DNA repair, and immune responses that warrants further investigation.

Our bulk RNA-seq and ATAC-seq analyses provide potential TFs involved in G1/S arrest mediated-expression of immune response in lymphoid cell lines **(Figure 5f)**. AP-1 (e.g., JUN, FOS, and ATF), IRF, and NF-κB TF families regulate immune response (Wagner and Eferl 2005, Oeckinghaus and Ghosh 2009, Ikushima, Negishi et al. 2013). In addition to regulating immune response, many of these TFs are also implicated in DNA damage response (Janssens and Tschopp 2006). Among them, *ATF3*, a TF from the AP-1 family, showed the strongest expression induction upon G1/S arrest **(Figure 5f)** and is known to regulate G1 to S transition (Liang, Wolfgang et al. 1996, Fan, Jin et al. 2002, Ku and Cheng 2020, Wu, Nicoll et al. 2021). We believe *ATF3* may play a role in the induction of IFN upon G1/S arrest in B and T cell lines and this induction may be DNA damage mediated. Taken together, our findings show the G1/S boundary as a cell intrinsic and independent regulator of IFN expression and immune response activation, and identify potential TFs orchestrating this response.

## Acknowledgements

We would like to thank Dr. Michael Zhang and Alyssa Briggs for their valuable feedback on the manuscript. We would also like to thank the Genome Center and Flow Cytometry core facilities at The University of Texas at Dallas for their support during the course of this research.

Author contributions: THK conceived, guided, and managed the project. GB designed, performed, and analyzed the experiments; GB performed all the wet lab experiments except for those provided as a service by UTD Genome Center which were bulk RNA-seq and scRNA-seq library preparation, QC, and sequencing. GB performed all the data analyses including construction of computational pipelines and custom scripts for the project. GB drafted the manuscript, and GB and THK together edited and revised the manuscript.

## Materials and Methods

### Cell Culture and Treatments

GM12878 and Ramos were obtained from Coriell Institute for Medical Research. Jurkat, THP-1, and IMR90 cells were purchased from American Type Culture Collection (ATCC). The cells were maintained according to the supplier’s instructions. 0.4 M/mL cells were seeded and incubated for 24h at 37 °C supplemented with 5% CO_2_ prior to treatment. Cells were treated with aphidicolin (Sigma-Aldrich) and nocodazole (Sigma-Aldrich) drugs to arrest them at G1/S phase and G2/M phase respectively at various time points. Sendai Virus (Cantrell strain) obtained from Charles River was used for inducing antiviral immune response as previously indicated (Banerjee, Kim et al. 2014).

GM12878 and Jurkat cells were treated in 4 conditions: Asynchronous (DMSO treated) and uninfected (CU), APH treated and uninfected (AU), DMSO treated and infected (CI), and APH treated and infected (AI). For bulk RNA-seq, both Jurkat and GM12878 treatments were performed in triplicates (24 conditions in total). For scRNA-seq DMSO and APH treated and infected GM12878 cells were used. For ATAC-seq, in GM12878, DMSO treatment was performed in triplicates whereas four replicates were used for the APH treatment (14 conditions in total). The DMSO/APH (1mg/mL) treatment was for 24h and the last 12h of this incubation was with SeV/no infection for infected samples and for controls respectively.

### Flow cytometry

To check the cell cycle phase, after the drug and SeV treatment, cells were harvested and stained with hypotonic propidium iodide (PI) stain (Riccardi and Nicoletti 2006). To remove RNA, Ambion RNase cocktail (Thermo Fisher Scientific) was used according to manufacturer’s instructions. Then the cells were strained and analyzed by BD LSRFortessa™ Flow Cytometer and FlowJo software.

### FACS

For ATAC-seq, cells death was measured with PE Annexin-V (BD Biosciences) according to manufacturer’s instructions. Cells were stained with 7-AAD viability staining solution (BD Biosciences) and 60,000 live cells per condition were sorted by BDAria Fusion and FlowJo software.

### ATAC-seq library preparation

Protocol was taken from different studies (Buenrostro, Giresi et al. 2013, Buenrostro, Wu et al. 2015, Corces, Trevino et al. 2017, Brunton, Garner et al. 2020) and solutions were prepared according to (Corces, Trevino et al. 2017). Briefly, 60K cells were sorted and lysed in 50 μL cold lysis buffer on ice for 3 min. Lysis was quenched by adding 1 mL of wash buffer. Cells were centrifuged at 500xg for 10 min at 4 °C and supernatant (cytoplasm) was discarded. Nuclei were gently resuspended with transposition mix and incubated at 37 °C for 30 minutes at 1,000 rpm. DNA was purified using DNA Clean & Concentrator-5 (Zymo Research, D4013) according to the manufacturer’s instructions. PCR amplification (library generation) was performed according to the previous study (Brunton, Garner et al. 2020) (Indexes used are in **Supplementary Table 8**). Library was purified using AMPure XP beads (Beckman Coulter) according to manufacturer’s instructions. To assess the library quality, each sample was run on an Agilent High Sensitivity DNA Bioanalysis chip following manufacturer’s instructions. DNA concentrations were measured by Qubit™ dsDNA HS Assay Kit (Invitrogen, Catalog Numbers Q32851, Q32854) following manufacturer’s instructions. 50 bp paired-end sequencing was performed (P3 flow cell by Illumina Nextseq 2000).

### ATAC-seq data analysis

The adapter removal was performed using cutadapt tool. Reads were aligned to hg38 genome using bwa-mem (Li and Durbin 2009). Quality control for the reads was performed using ATACQC (Ou, Liu et al. 2018). PCR duplicates, unpaired reads, mitochondrial reads, and bad quality reads (reads with MAPQ score less than 30) were removed using bamtools (Barnett, Garrison et al. 2011). Bam files per condition (CU, CI, AU, and AI) were pooled and peaks were called using macs2 with the following parameters: callpeak --cutoff-analysis -f BED --shift -75 --extsize 150 -- nomodel --keep-dup all -p 0.01 (Zhang, Liu et al. 2008). Peaks were called for each sample (14 samples) with macs2 using the same parameters. The intersection of peaks with the pooled peaks and the peaks per sample were extracted using bedtools intersect command and each peak was given a unique name (Quinlan and Hall 2010). To get the common peaks, all intersected peaks per condition were merged with bedtools merge and count matrix was created with FeatureCounts (Quinlan and Hall 2010, Liao, Smyth et al. 2014). Differential accessibility analysis was performed using DESeq2 (Love, Huber et al. 2014) between four conditions (CU, CI, AU, and AI). Differential accessibility region annotation was performed using GREAT (McLean, Bristor et al. 2010). Motif analysis within these differentially accessible regions were performed with HOMER (Heinz, Benner et al. 2010).

### RNA Extraction, DNase Treatment, cDNA Synthesis, and Real Time Quantitative PCR

RNA was extracted using Qiagen RNeasy kit according to the manufacturer’s instructions. Extracted RNA was treated with DNase by TURBO DNase (Thermo Fisher Scientific) to remove any genomic DNA contamination according to the manufacturer’s instructions. The DNase was removed using RNA Clean & Concentrator-25 kit according to the supplier’s instructions (Zymo Research). Reverse transcription was done using cDNA ImProm-II reverse transcription kit (Promega) according to the manufacturer’s instructions. RT-qPCR was performed using FastStart Universal SYBR Green Master (Sigma-Aldrich) and primers are given in **Supplementary Table 9**.

### Bulk RNA-seq library preparation

RNA-seq library preparation was performed using TruSeq Stranded mRNA kit (Illumina) according to the manufacturer’s instructions. Sequencing was performed using Illumina Nextseq 2000.

### Bulk RNA-seq data analysis

The reads were aligned to hg38 genome and the reads with proper pairs and MAPQ score > 30 were selected for downstream analysis. Count matrix was prepared using FeatureCounts (Liao, Smyth et al. 2014). Differential gene expression analysis was performed using DESeq2 (Love, Huber et al. 2014) between four conditions (CU, CI, AU, and AI). Gene ontology enrichment analysis within these differentially expressed genes were performed with clusterProfiler (Yu, Wang et al. 2012).

### scRNA-seq library preparation

scRNA-seq library preparation was performed using 10x Genomics Chromium 3’ single cell library protocol (Illumina) according to the manufacturer’s instructions. Sequencing was performed using Illumina Nextseq 2000.

### scRNA-seq data analysis

The reads were aligned to hg38 genome and filtered using default CellRanger 3.1.0 command (Zheng, Terry et al. 2017). Dimensionality reduction, clustering of cells, and cell cycle score calculations were performed using Seurat v4 (Hao, Hao et al. 2021). Batch correction was performed with Harmony (Korsunsky, Millard et al. 2019). Pseudotime analysis based on cell cycle phases was performed using Destiny (Haghverdi, Buttner et al. 2016).

## Supplementary Data

**Supplementary Figure 1:**
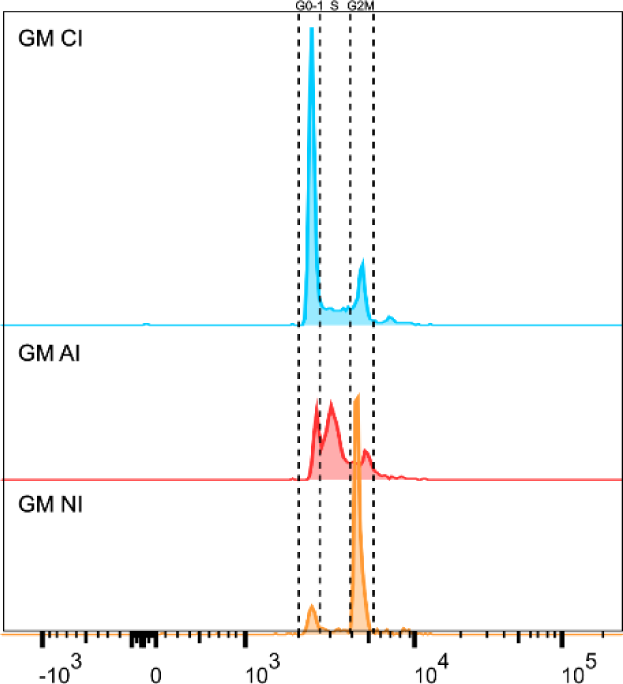
DNA content histograms after propidium iodide staining to determine the cell cycle stages of SeV infected GM12878 cells were treated with: DMSO as control (GM CI), APH (GM AI), and NOCO (GM NI). Cell cycle phases are indicated above the histrograms.

**Supplemental Figure 2:**
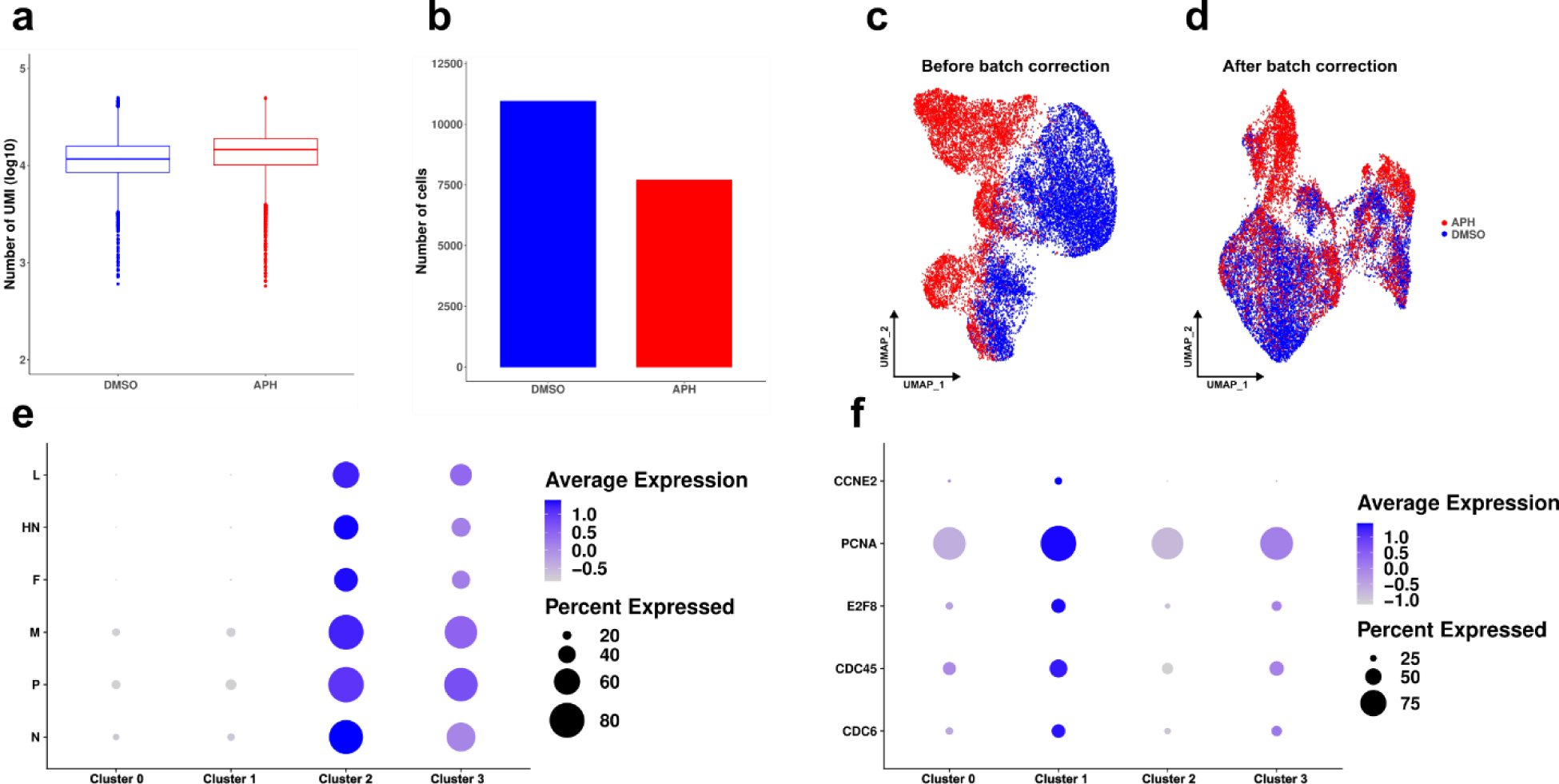
Additional analyses of scRNA-seq data in GM12878 (GM) cells across treatment conditions. **(a)** Number of UMIs in log scale in DMSO treated GM and APH treated GM samples (x-axis). Blue indicates DMSO treated GM and red indicates APH treated GM. **(b)** Bar plot of number of cells across treatments. **(c)** UMAP of cells before batch correction and **(d)** after batch correction. **(e)** Bubble plot showing the expression of viral genes across clusters of cells. **(f)** Bubble plot showing the expression of S phase genes across clusters of cells.

**Supplementary Figure 3:**
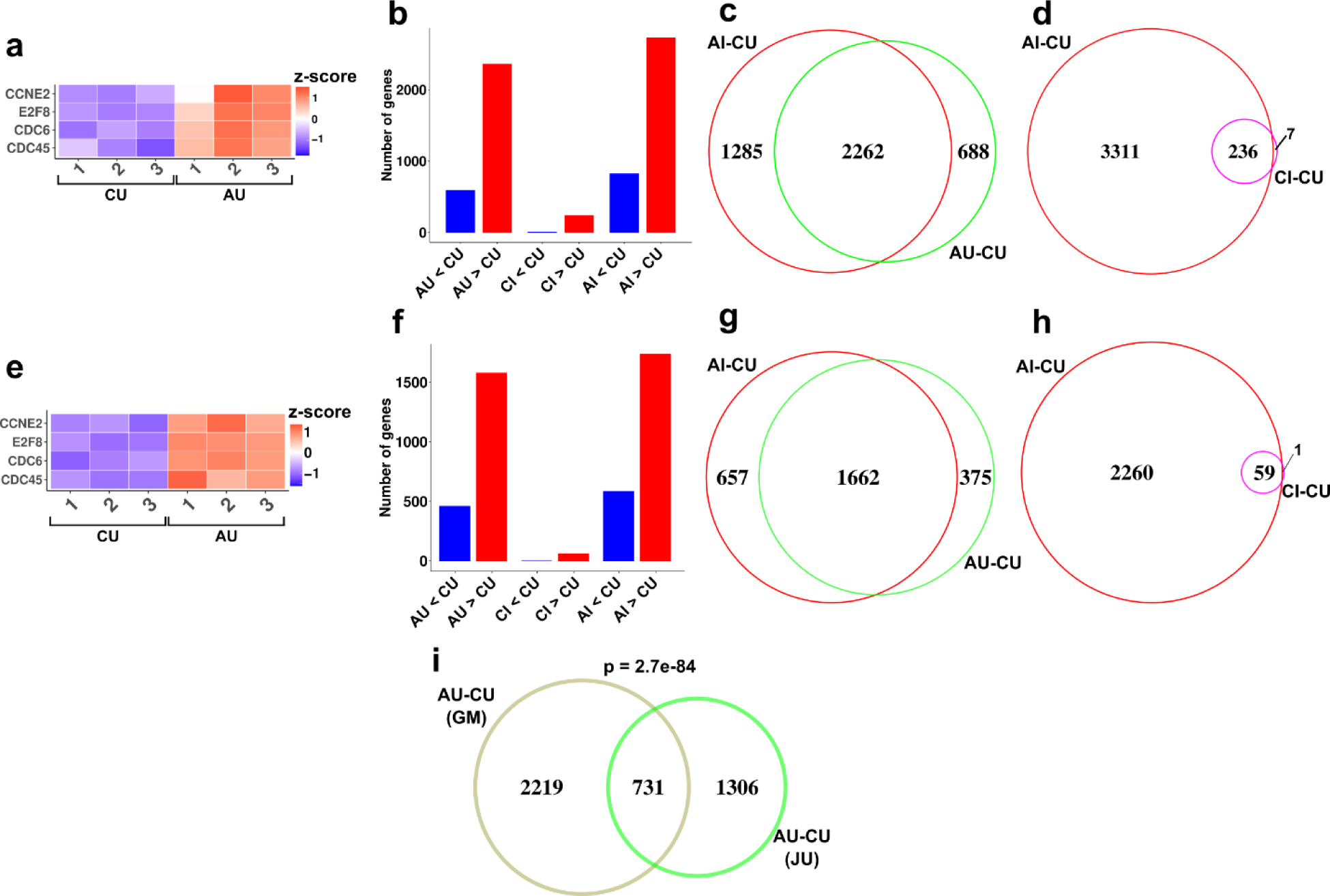
Additional analyses of RNA-seq data in GM12878 (GM) and Jurkat (JU) cell lines across treatment conditions. **(a)** Heatmap of S-phase marker genes in AU compared to CU samples in GM cells. CU: control uninfected, AU: arrested uninfected, CI: control infected, AI: arrested infected. Numbers denote biological replicates. Gene expression values were normalized and z-transformed across all samples. **(b)** Number of differentially expressed genes in pairwise comparisons in GM cells (x-axis). Blue indicates downregulation and red indicates upregulation compared to the baseline. **(c)** Venn diagram of gene overlap between AI-CU and AU-CU comparisons in GM cells. A gene was considered overlapping if it was differentially expressed in the same direction between the two comparisons. **(d)** Same as (c), but between AI-CU and CI-CU. **(e-h)** Same as a-d but for JU cells. **(i)** Overlap of differentially expressed genes between AU-CU in GM and JU cells. Statistical significance was determined by Fisher’s exact test.

**Supplementary Figure 4:**
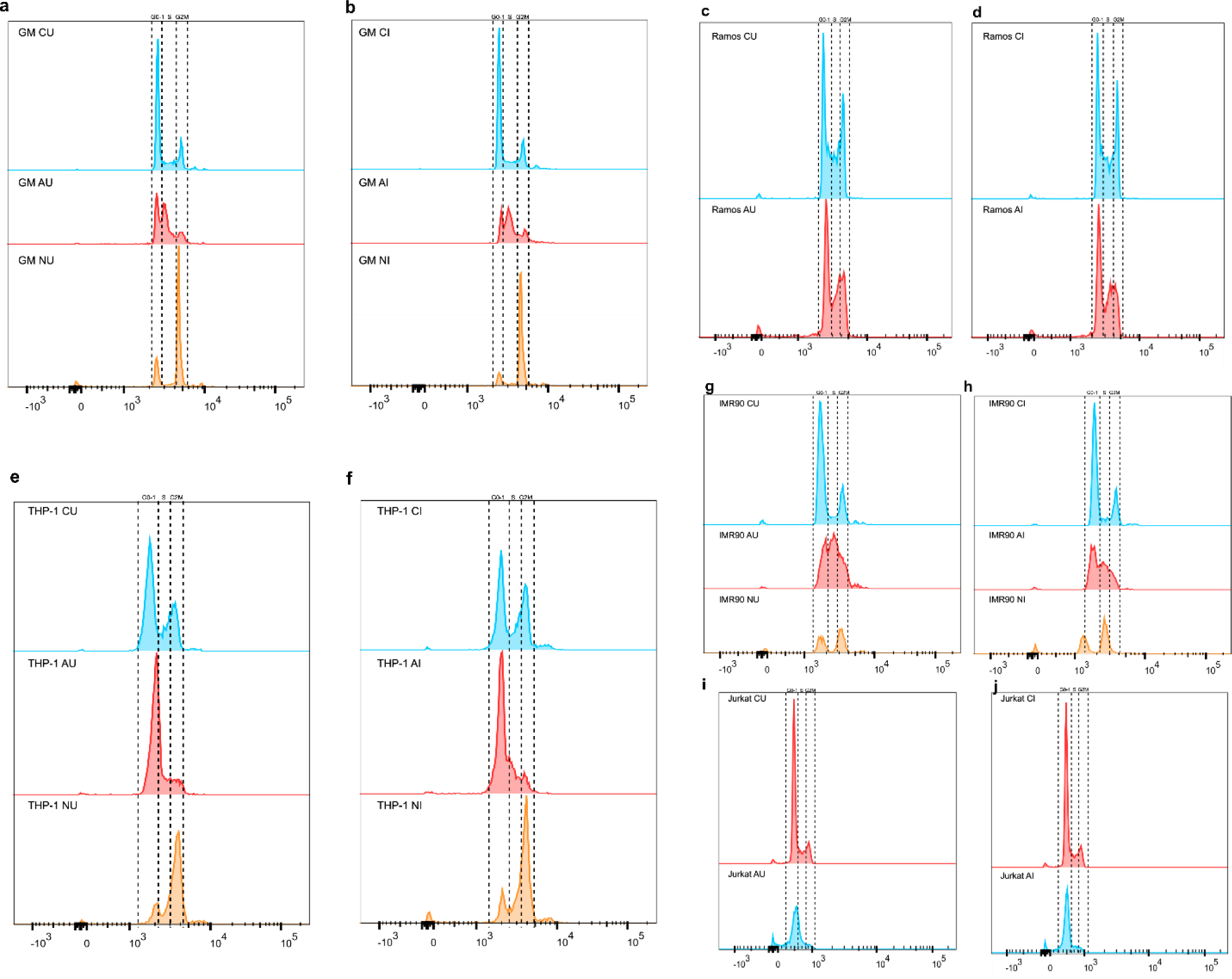
DNA content histograms after propidium iodide staining to determine the cell cycle phases across treatment conditions with DMSO as control (CU), aphidicolin (AU), nocodazole (NU), SeV infection-DMSO (CI), SeV infection-aphidicolin (AI), and SeV infection-nocodazole (NI). These treatments were performed for: **(a-b)** GM12878, **(c-d)** Ramos, **(e-f)** THP-1, **(g-h)** IMR90, and **(i-j)** Jurkat. Cell cycle phases are indicated above the histograms.

**Supplemental Figure 5:**
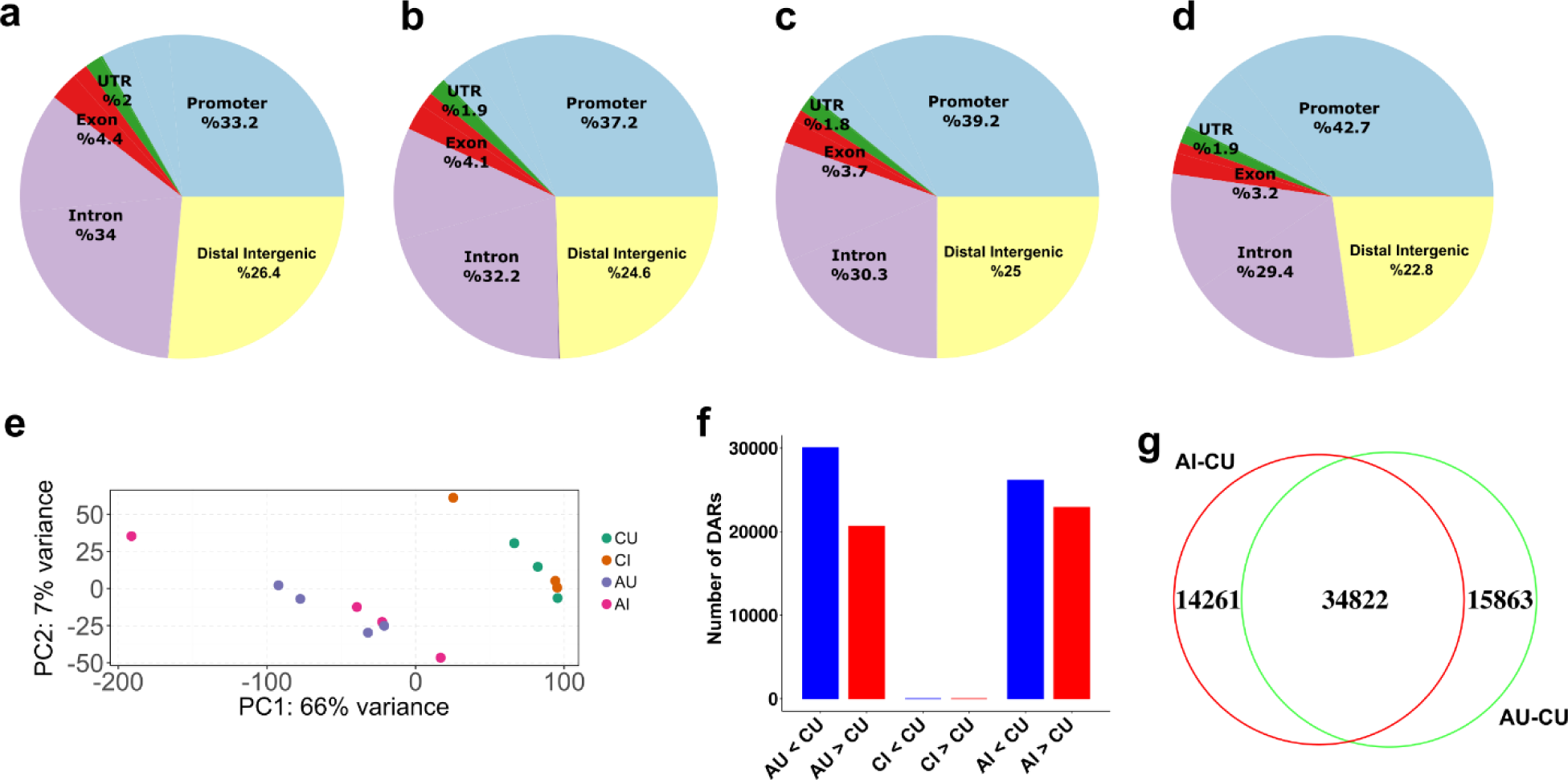
Additional analyses of ATAC-seq data in GM12878 (GM) cell line across treatment conditions. **(a-d)** Breakdown of peak annotation across genomic regions per condition (left to right: CU, CI, AU, AI). **(e)** Principal component analysis of peak count data across biological samples. Note that PC1 and PC2 separates arrested and control samples. **(f)** Number of differentially accessible regions (DARs) in pairwise comparisons in GM cells (x-axis). Blue indicates closed regions and red indicates open regions compared to the baseline. CU: control uninfected, AU: arrested uninfected, CI: control infected, AI: arrested infected. (g) Venn diagram of differentially accessible region overlap between AU-CU and AU-CU comparisons in GM cells. A region was considered overlapping if it was differentially accessible in the same direction between the two comparisons.

**Supplementary Table 1: List of differentially expressed genes in bulk RNA-seq across conditions in GM12878 cells.**

**Supplementary Table 2: List of gene ontology enrichment of differentially expressed genes in bulk RNA-seq across conditions in GM12878 cells.**

**Supplementary Table 3: List of differentially expressed genes in bulk RNA-seq across conditions in Jurkat cells.**

**Supplementary Table 4: List of gene ontology enrichment of differentially expressed genes in bulk RNA-seq across conditions in Jurkat cells.**

**Supplementary Table 5: List of differentially accessible regions in ATAC-seq across conditions.**

**Supplementary Table 6: List of gene ontology enrichment of differentially accessible regions in ATAC-seq across conditions.**

**Supplementary Table 7:**
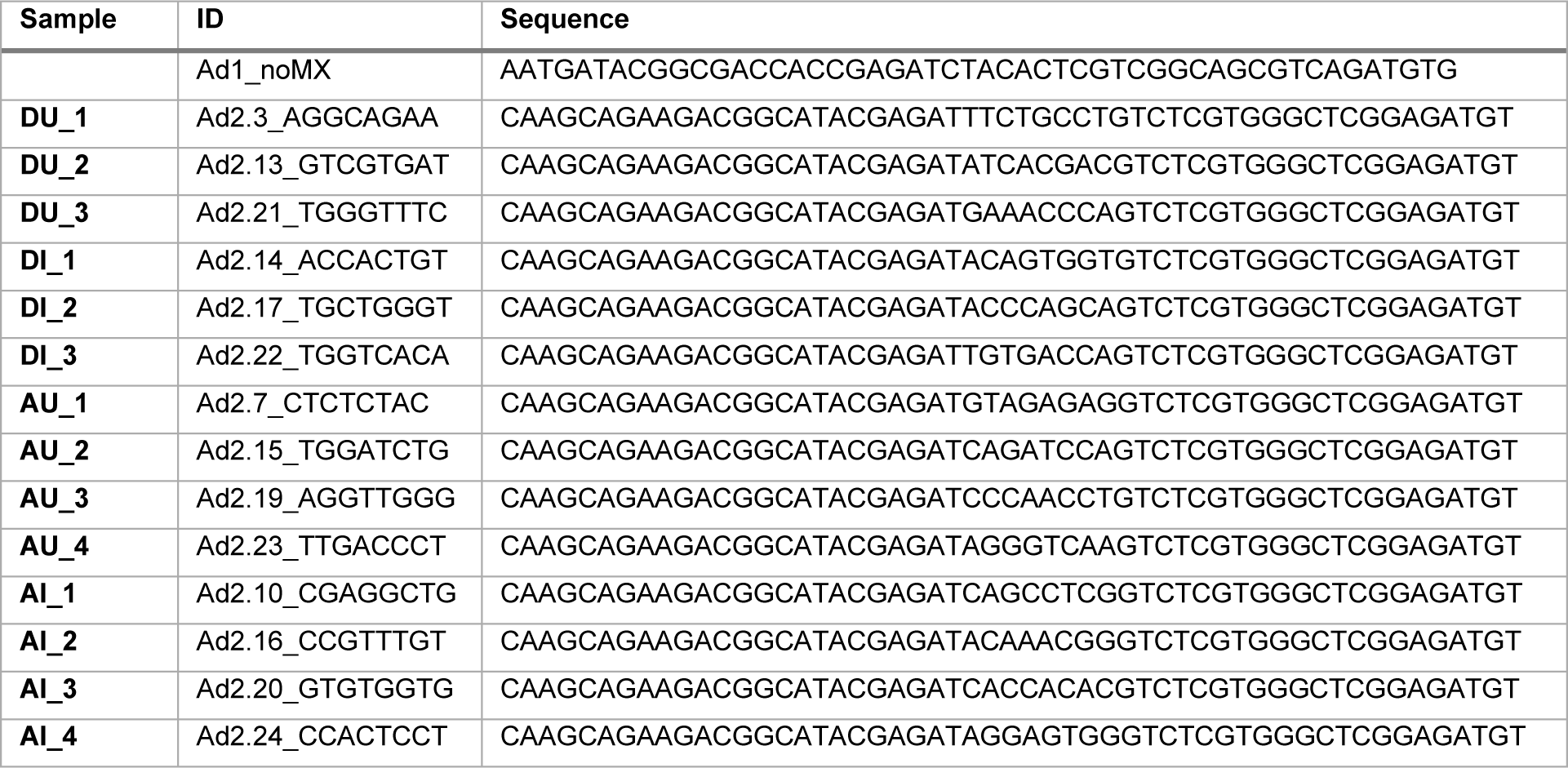
List of motifs which were significantly enriched within differentially open regions in AU compared to CU.

**Supplementary Table 8:**
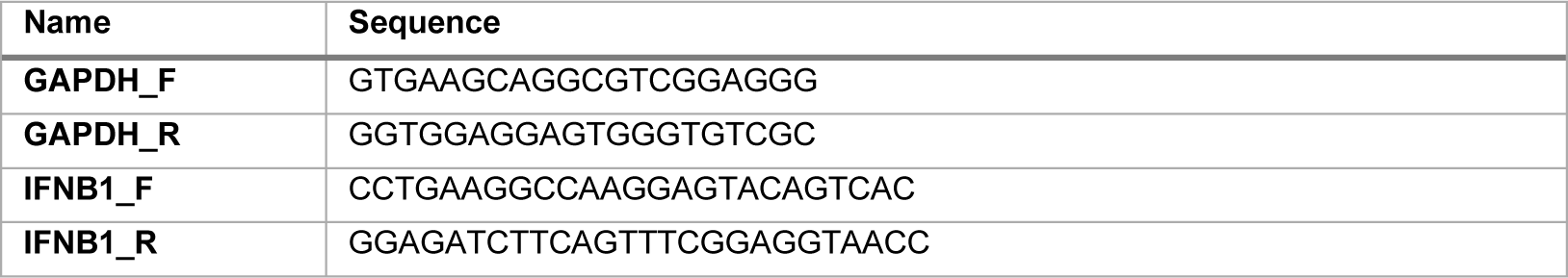
Illumina/Nextera i5 common adapter and i7 index adapters for ATAC-seq.

**Supplementary Table 9: Primers used in RT-qPCR**

## Notes

### Competing Interest Statement

The authors have declared no competing interest.

### Summary of Updates

Edits on author contributions.

